# The omnitig framework can improve genome assembly contiguity in practice

**DOI:** 10.1101/2023.01.30.526175

**Authors:** Sebastian Schmidt, Santeri Toivonen, Paul Medvedev, Alexandru I. Tomescu

**Author notes:** Shared last-author contribution (Paul Medvedev, Alexandru I. Tomescu).

## Abstract

Despite the long history of genome assembly research, there remains a large gap between the theoretical and practical work. There is practical software with little theoretical underpinning of accuracy on one hand and theoretical algorithms which have not been adopted in practice on the other. In this paper we attempt to bridge the gap between theory and practice by showing how the theoretical safe-and-complete framework can be integrated into existing assemblers in order to improve contiguity. The optimal algorithm in this framework, called the omnitig algorithm, has not been used in practice due to its complexity and its lack of robustness to real data. Instead, we pursue a simplified notion of omnitigs, giving an efficient algorithm to compute them and demonstrating their safety under certain conditions. We modify two assemblers (wtdbg2 and Flye) by replacing their unitig algorithm with the simple omnitig algorithm. We test our modifications using real HiFi data from the Drosophilia melanogaster and the Caenorhabditis elegans genome. Our modified algorithms lead to a substantial improvement in alignment-based contiguity, with negligible computational costs and either no or a small increase in the number of misassemblies.

## 1 Introduction

Genome assembly is a classical problem in bioinformatics that has received a lot of both theoretical and practical attention. On the practical side, many successful assemblers have been developed and have led to numerous biological discoveries (e.g. [Nur+22; Rhi+21]). On the theoretical side, there have been many attempts at modelling the problem and coming up with algorithms whose accuracy is optimal with respect to these models (e.g. [TM17; OMT18; Cai+19; Cai+21; Cai+20; Cai+22]). Unfortunately, there remains a large gap between the theoretical and practical work, resulting in practical software with little theoretical underpinning of accuracy on the one hand and theoretical algorithms which have not been adopted in practice on the other [Med22].

The first group of theoretical approaches formulated assembly as the problem of finding a complete reconstruction of the genome (i.e. one string per genome). Initially, the proposed algorithm was to find a Eulerian cycle in a genome graph [PTW01]. Later work described more sophisticated algorithms that maximised the probability of successfully finding the complete reconstruction, if one exists [BBT13]. However, these formulations did not capture the fact that the conditions under which a complete reconstruction is feasible are extremely rare [BBT13; MP21]. Therefore, the second group of theoretical approaches formulated assembly as the problem of finding a set of sequences (called *omnitigs*) that were as long as possible and were guaranteed to be substrings of the genome (i.e. *safe*) [TM17]. It was possible to formally characterise how omnitigs looked like in the genome graph and how to efficiently find all possible omnitigs (i.e. a *complete* algorithm) in an idealised setting [Cai+19; Cai+21; Cai+20]. However, the omnitig algorithm requires complex data structures [Cai+21; GIP20] and omnitigs themselves are not safe in the presence of sequencing errors, missing coverage, or linear chromosomes [TM17]; as a result, omnitigs have not been applied in practice. Instead, most assembly software use the much simpler and more accurate unitig algorithm.

In this paper we attempt to bridge the gap between theory and practice by showing how the safe-and-complete framework can be integrated into existing assemblers in order to improve contiguity. To do so, we pursue a relaxed notion of omnitigs, called *simple omnitigs*. Simple omnitigs are walks having a non-branching *core*, such that all nodes to the right of the core have out-degree one (i.e., a unique right extension), and all nodes to the left of the core have in-degree one (i.e., a unique left extension) (see Figure 1(a) for an illustration). The idea behind simple omnitigs was in fact known in the literature also before the safety framework (e.g. [Med+07; Jac09; KSP10]), and also called Y-to-V transformation [TM17]. Though simple omnitigs are known to be not complete, they are nevertheless safe for a single circular chromosome [TM17]. On perfect data, they were shown to significantly improve length and contiguity over unitigs, while almost reaching that of omnitigs [TM17]. We therefore consider simple omnitigs as the “compromise candidate” for adopting the safe-and-complete framework to be used in practice.

**Fig. 1:**
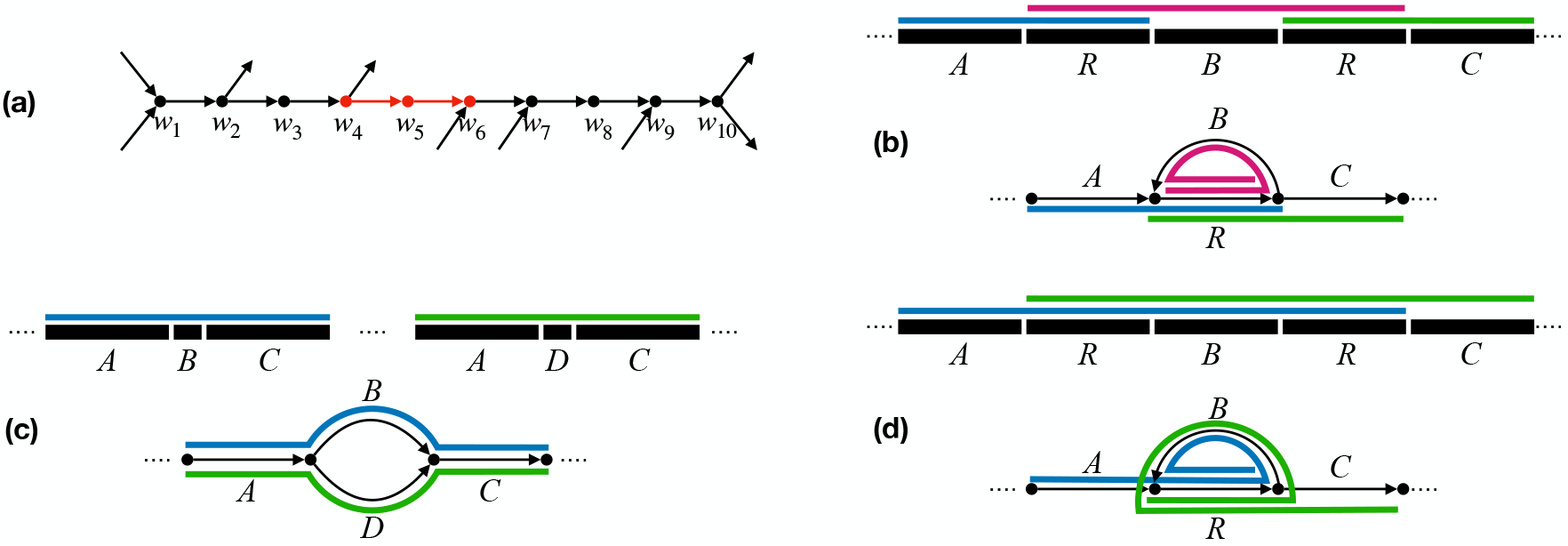
Overview of simple omnitigs. In (a), we show a simple omnitig, with its core highlighted in red. By definition, the core is a unitig (only the first and last node are branching). Since the walk is a simple omnitig, nodes *w*_2_, *w*_3_, *w*_4_ have exactly one incoming arc, and nodes *w*_6_, *w*_7_, *w*_8_, *w*_9_ have exactly one outgoing arc. The inner nodes of the core (i.e., *w*_5_) have exactly one incoming and exactly one outgoing arc. In (b) and (c), on top, we show the repeat structure at genome level, where we assume each labelled substring corresponds to a unitig; on the bottom, we show the repeat structure at assembly graph level, where we draw each unitig as a length-two path. Simple omnitigs are shown as coloured lines. In (b), there are two occurrences of a repeat *R*, with simple omnitigs providing more context to *R*, in this case even capturing the full substring from the first to the second occurrence of *R*. In (c), a substring *ABC* also appears in the variant *ADC* (with *B* replaced by *D*), which at the graph level induces a bubble. Simple omnitigs provide context around the variants *B* and *D*, in this case by recovering the full strings *ABC* and *ADC*. In (d), we show the omnitigs of the genome graph of (b) for comparison. Omnitigs allow to safely extend the repeat *R* both forwards and backwards within a single walk (*ARBR* and *RBRC*). The omnitigs in (c) are equivalent to simple omnitigs.

In this paper, we prove that simple omnitigs remain safe even when there are multiple linear chromosomes, as long as no chromosome starts or ends inside them. We give a linear output-sensitive time algorithm for finding all simple omnitigs. We then integrate simple omnitigs into two widely-known graph-based genome assemblers, Flye [Kol+19] and wtdbg2 [RL20]. We use real HiFi data to demonstrate that the integration substantially improves assembly contiguity with negligible computational cost. On *D. melanogaster*, such assemblies have the same level of correctness, while having substantially higher alignment-based contiguity metrics than the original assemblers. Our extension of wtdbg2 results in similar contiguity and accuracy as the most accurate assembler on this dataset (HiCanu [Nur+20]) but is more than 20 times faster. On *C. elegans*, we improve the alignment-based contiguity of the best performing assembler Flye, albeit with a small increase in misassembly errors. We also improve the contiguity of wtdbg2, this time without any additional errors.

## 2 Results

### 2.1 Simple omnitigs

Many assemblers work by constructing some kind of genome graph and outputting unitigs from it. A unitig is a path whose inner nodes have in- and out-degree one. Instead, we propose outputting *simple omnitigs*.

A simple omnitig is a walk in a graph, defined as the *univocal extension* of a unitig. The univocal extension is the maximal superwalk of the unitig that contains only inner nodes with in-degree one, including the first node of the unitig, then the unitig, and then only inner nodes with out-degree one, including the last node of the unitig. The unitig that “produces” a simple omnitig in this way is called a *core* (note that simple omnitigs may contain multiple unitigs, but only one of them can be the core). Figure 1 (a) shows an example of a unitig and a simple omnitig. In Section 3.2, we will prove that, under idealised assumptions, simple omnitigs in the de Bruijn graph of the reads are mostly guaranteed to be correct (Theorem 1). Then, in Section 3.3, we show how the core is a useful algorithmic building block (Theorem 2) and give an algorithm to output all simple omnitigs in linear time.

Simple omnitigs are longer than unitigs and thus provide more context in the final assembly. They are effective for example in repeats or bubbles, as the contigs do not need to stop at the start or end of the structure, but can continue through the flanks as well. See Figure 1 (b) and (c) for an example, and also Figure 1 (d) for a comparison to omnitigs. Besides these examples, simple omnitigs also increase contig length in tangled regions of the graph that contain more complex structures than bubbles and repeats. Whenever there is a node with indegree 1 but larger outdegree (or vice-versa), simple omnitigs can connect the unitigs starting or ending in such a node to produce longer contigs.

### 2.2 Injecting simple omnitigs into existing assemblers

In order for an assembler to be modifiable to output simple omnitigs instead of unitigs, it needs to work by building some kind of assembly graph and outputting unitigs from it. Furthermore, if the assembler does additional processing of the unitigs, then this processing needs to be either disabled or modified so that it becomes compatible with simple omnitigs. We identified two assemblers that lent themselves to the integration of simple omnitigs. Full details are in Appendix A, but we summarise the changes here.

The first assembler is wtdbg2 [RL20], which builds a “fuzzy de Bruijn graph” and then uses unitigs from this graph to build a “fragment graph.” It then performs further error corrections on the fragment graph before finally outputting the consensus sequences of its unitigs as contigs. We made two modifications. First, we changed the fuzzy de Bruin graph construction so that it takes advantage of homopolymer compressed space. This was needed to improve the underlying quality of the graph, which is especially important for simple omnitigs. Second, we changed the final output to be the consensus of simple omnitigs (rather than of unitigs) on the error-corrected fragment graph.

The second assembler is Flye [Kol+19], which constructs a “repeat graph” followed by a repeat resolving and polishing step prior to outputting unitigs. We modified Flye by 1) disabling the post construction step of repeat resolving and polishing, and 2) outputting simple omnitigs instead of unitigs. We disabled the resolving and polishing steps because they were incompatible with injecting simple omnitigs. To make sure that this did not hamper Flye, we verified that this change did not significantly alter the contiguity of the assembly by running it with only the first modification.

### 2.3 Experimental setup

To evaluate the performance of our modified assemblers, we use two real datasets of Pacbio HiFi reads, one from D. melanogaster and the other from C. elegans. We use all chromosomes from each dataset for evaluation. Table 1 shows the properties of these reference genomes and the corresponding reads. We measure accuracy and contiguity using a modified QUAST-LG [Mik+18] tool and the reference genome of the same D. melanogaster individual (`GCF 000001215.4`) and a different C. elegans individual (`GCA 000002985.3`). Following the observations of [Ban+22], we modified QUAST-LG to work in homopolymer compressed space. Otherwise, as [Ban+22] observed, QUAST-LG falsely reports misassemblies on genomic regions with long homopolymer runs.

**Table 1:**
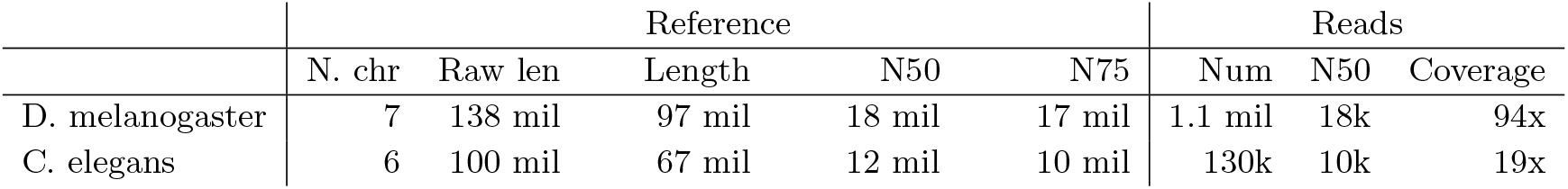
Properties of the reference genomes and the HiFi reads we use. All numbers (except “Raw len” and “Coverage”) are shown in homopolymer compressed space. Coverage is given with respect to the diploid genome length.

|  | Reference |  |  |  |  | Reads |  |  |
| --- | --- | --- | --- | --- | --- | --- | --- | --- |
|  | N. chr | Raw len | Length | N50 | N75 | Num | N50 | Coverage |
| D. melanogaster | 7 | 138 mil | 97 mil | 18 mil | 17 mil | 1.1 mil | 18k | 94x |
| C. elegans | 6 | 100 mil | 67 mil | 12 mil | 10 mil | 130k | 10k | 19x |

Unlike unitigs, simple omnitigs can overlap, resulting in the same reference sequence being potentially present in more than one simple omnitig. This makes some QUAST-LG metrics misleading or inappropriate. In particular, we did not use the NGA50/NGA75 contiguity metrics that QUAST-LG reports by default. Instead, we implemented and used the EA50max and EA75max metrics, which are similar but robust against overlapping contigs. These work by aligning the contigs against the reference, identifying for each reference position the longest alignment (i.e. to a contig or part of a contig, if the contig is misassembled), and then reporting the 50th and 75th percentiles of the distribution of these lengths. For example, an EA50max value of *ℓ* means that 50% of the genomic positions are covered by a contig (or part of a contig, if misassembled) of length at least *ℓ*. The choice for the longest alignment was made because otherwise assemblers would be penalised for overlaps between longer and shorter contigs, even though the existence of the shorter contig does not reduce the quality of the longer contig (see Section 3.4 for more details). Furthermore, we modified QUAST-LG to only report at most one misassembly per reference position. Otherwise, a single misassembly in a unitig will count multiple times if there are numerous simple omnitigs containing that unitig. Section 3.4 describes these modifications, as well as their justifications, in more detail.

In order to establish a baseline of the state-of-the-art assembly performance on our datasets, we ran hifiasm [Che+21], LJA [Ban+22] and HiCanu [Nur+20]. All were run with default parameters (using the predefined mode for HiFi reads in HiCanu).

### 2.4 D. melanogaster

Figure 2a shows that simple omnitigs lead to a substantial improvement in assembly contiguity. For wtdbg2, this especially happens for the shorter contigs. For example, the EA75max is increased to 9.9 mil from 4.0 mil when incorporating simple omnitigs. This is consistent with previous observations on error-free data [TM17] and due to the fact that simple omnitigs typically extend contigs through parts of the graph that are more tangled and hence contain shorter unitigs. For Flye, EA*x*max is increased across the board, however the quality of the Flye assembly pipeline is low on this dataset, with or without modifications.

**Fig. 2:**
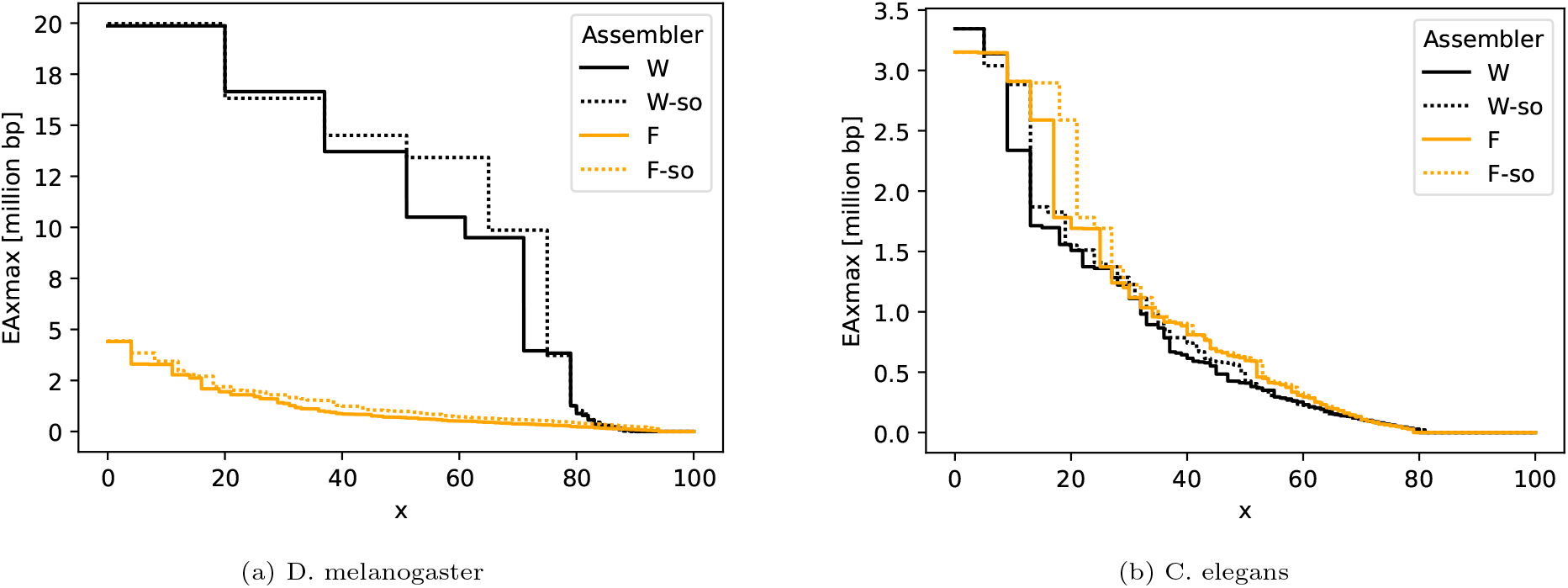
Assembly contiguity of wtdbg2 (denoted by “W”) and Flye (denoted by “F”) and their simple omnitig versions (denoted with a “-so” suffix). The y-axis shows the EA*x*max metric, for 0 ≤ *x* ≤ 100. Differences between the curves on smaller *x* pertain changes in longer contigs, and for larger *x* they pertain changes in shorter contigs (same as with the well-known NGA*x* metric family).

Table 2 shows the full statistics for D. melanogaster assemblies, including for other assemblers. Overall, compared to the results of three other assemblers on this data, our modified wtdbg2 pipeline achieves the highest contiguity on longer contigs (i.e. an EA50max of 14.5 MB), at the cost of three more misassemblies. We note that the contiguity metrics take the misassemblies into account, i.e. the length statistics are calculated after breaking contigs apart at the misassemblies.

**Table 2:** Assembly accuracy on D. melanogaster. All statistics are shown in homopolymer compressed space. “W-int” is wtdbg2 with the homopolymer modification but without simple omnitigs, and “F-int” is Flye with the postconstruction steps disabled but without simple omnitigs. ^*†*^: Note that this number is much worse than the NGA50 reported for D. melanogaster on https://github.com/fenderglass/Flye. This is likely due to us using the highly heterozygous cross of the A4 and ISO1 strains of D. melanogaster, while the variant assembled on the Flye website is plain ISO1.

|  | wtdbg2 |  |  | Flye |  |  | Others |  |  |
| --- | --- | --- | --- | --- | --- | --- | --- | --- | --- |
|  | W | W-int | W-so | F | F-int | F-so | hifiasm | LJA | HiCanu |
| EA50max ($\times 10^6$) | 13.7 | 12.2 | 14.5 | 0.7<sup>†</sup> | 0.7 | 1.0 | 14.1 | 6.0 | 13.4 |
| EA75max ($\times 10^6$) | 4.0 | 3.8 | 9.9 | 0.3 | 0.3 | 0.5 | 4.8 | 1.7 | 13.0 |
| N. misassemblies | 1 | 1 | 4 | 3 | 3 | 5 | 1 | 1 | 0 |
| N. contigs | 405 | 369 | 363 | 856 | 951 | 576 | 1,871 | 925 | 1,110 |
| Genome fraction (%) | 90.1 | 89.2 | 89.1 | 94.9 | 94.9 | 94.7 | 95.7 | 95.5 | 95.3 |
| Duplication ratio | 1.02 | 1.01 | 1.55 | 1.68 | 1.68 | 1.98 | 1.99 | 1.74 | 1.81 |

We also confirmed that these improvements are due to simple omnitigs and not to the other modifications we made. Table 2 shows the EA50max and EA75max numbers for the intermediate versions of these assemblers which contain all the non-simple omnitig modifications. Their contiguity does not improve relative to the original assemblers.

Table 2 also shows other assembly statistics. Notably, there is a small increase in the number of misassemblies. We believe that this stems from errors of the sequences stored at the branching nodes (i.e. with an in- or out-degree of more than one) in the graph. In particular, such nodes always lie at the ends of unitigs (by definition of unitig). An artefact of QUAST-LG is that errors in the last ≈ 1kbp of a contig are not counted as a misassembly, hence errors in most branching nodes do not affect the number of misassembled unitigs. However, branching nodes are often absorbed into the middle of simple omnitigs, causing QUAST-LG to report a misassembly error. Another effect of using simple omnitigs is that there is no longer a unique contig for each region of the genome. Table 2 shows the duplication ratio, which is higher for simple omnitigs than for unitigs. This is expected and in fact desired, since a region can flank and provide context for multiple bubbles at the same time. Note that most assemblers report a duplication ratio higher than one, since D. melanogaster is diploid, but the reference contains only one copy of each chromosome.

Finally, our modification also have a slight affect on the genome fraction, but, since these are minor, we did not investigate these further. The major differences in genome fraction between the different baseline assemblers is due to inherent differences between those assemblers, rather than any effect of our modifications.

### 2.5 C. elegans

Figure 2b shows that simple omnitigs improve both wtdbg2 and Flye in contiguity for longer contigs (i.e. small values of *x*). In Table 3 we see that simple omnitigs increase the EA50max of wtdbg2 by 20% over the intermediate variant (and even more over default wtdbg2), with the same number of misassemblies as default wtdbg2. The improvement of Flye is mostly invisible in EA50max or EA75max, since it is only present within less than 50% of the reference length. Overall, comparing between all assemblers, Flye achieves the best EA50max among the baseline assemblers, and the simple omnitig variant improves this even further, albeit at a cost of three more misassemblies (recall, however, that the EA*x*max metrics are computed on the contigs after breaking them at all misassemblies).

**Table 3:** Assembly accuracy on C. elegans. All statistics are shown in homopolymer compressed space. Some of the assemblers achieve less than 75% genome fraction, hence their EA75max is undefined.

|  | wtdbg2 |  |  | Flye |  |  | Others |  |  |
| --- | --- | --- | --- | --- | --- | --- | --- | --- | --- |
|  | W | W-int | W-so | F | F-int | F-so | hifiasm | LJA | HiCanu |
| EA50max ($\times 10^3$) | 412 | 425 | 512 | 620 | 620 | 628 | 596 | 591 | 342 |
| EA75max ($\times 10^3$) | 68 | 66 | 67 | 64 | 63 | 63 | 43 | - | - |
| N. contigs | 389 | 394 | 392 | 280 | 276 | 274 | 346 | 239 | 475 |
| N. misassemblies | 4 | 2 | 4 | 2 | 2 | 5 | 6 | 1 | 0 |
| Genome fraction (%) | 80.8 | 80.8 | 81.2 | 79.8 | 79.8 | 79.8 | 77.4 | 72.4 | 74.8 |
| Duplication ratio | 1.00 | 1.00 | 1.02 | 1.00 | 1.00 | 1.05 | 1.01 | 1.01 | 1.01 |

Table 3 also show the genome fraction and duplication ratio metrics. As with D.melanogaster, our modifications have a negligible effect on the genome fraction, while increasing the duplication ratio. The increase is expected, since a single reference position can now be covered by more than one contig. We note that the genome fraction of all assemblers is only ≤ 81% and the duplication ratio is only ∼ 1, even though the genome is diploid. We suspect that these numbers may be due to the relatively low coverage of this dataset. In any case, the genome fraction and duplication ratios are a property of the dataset and the baseline assemblers, rather than our modifications; hence, we do not investigate them further.

### 2.6 Time and memory usage

Table 4 shows the time and memory usage of the assemblers. Our modification did not affect the memory usage for Flye and only slightly for wtdbg2. Adding simple omnitigs to Flye decreased its run-time, since we disabled a post processing step. For wtdbg2, the running time increases by 9-61%, though it remains very fast (e.g. is is 3−25 times faster than the next fastest assembler). Since the focus of our study was contiguity and accuracy, we did not optimise our code for speed, and it is likely that the time increase could be mostly avoided by removing superfluous disk I/O.

**Table 4:** Time and memory usage.

|  |  | wtdbg2 |  | Flye |  | Others |  |  |
| --- | --- | --- | --- | --- | --- | --- | --- | --- |
|  |  | W | W-so | F | F-so | hifiasm | LJA | HiCanu |
| time (s) | D. melanogaster | 942 | 1,521 | 24,465 | 20,261 | 75,936 | 63,307 | 39,118 |
|  | C. elegans | 208 | 226 | 1,269 | 830 | 858 | 695 | 769 |
| mem (GiB) | D. melanogaster | 17 | 15 | 119 | 119 | 105 | 68 | 19 |
|  | C. elegans | 4.5 | 5.7 | 17 | 17 | 18 | 6.6 | 5.2 |

## 3 Methods

In this section, we will give a more formal definition of simple omnitigs, a proof of their safety, and a fast algorithm for computing them. We also describe our approach to evaluation of assemblies.

### 3.1 Definitions

A *graph G* = (*V, E*) is defined to be directed with *n nodes* in *V* and *m arcs* in *E*. The *tail* of an arc *e* = (*u, v*) is tail(*e*) = *u* and its *head* is head(*e*) = *v*. A *w*_1_-*w*_*ℓ*_ *walk W* = (*w*_1_, …, *w*_*ℓ*_) is a sequence of adjacent nodes. Its *tail* is tail(*W*) = *w*_1_ and its *head* is head(*W*) = *w*_*ℓ*_. A graph is *strongly connected* if each pair of nodes is connected by a walk. The concatenation of two walks *W* = (*w*_1_, …, *w*_*ℓ*_) and *X* = (*x*_1_, …, *x*_*ℓ*_) where *w*_*ℓ*_ = *x*_1_ is a walk *WX* = (*w*_1_, …, *w*_*ℓ*_, *x*_2_, …, *x*_*ℓ*_). A *split* is a node with at least two outgoing arcs and a *join* is a node with at least two incoming arcs. Let *W* = (*w*_1_, …, *w*_*ℓ*_) be a walk with *ℓ ≥* 2. The *inner* nodes of *W* are *w*_2_, …, *w*_*ℓ−*1_. A *unitig* is a walk whose inner nodes have in-and out-degree one ^1^. Let *w*_*i*_ be its first inner join, or *w*_*ℓ*_ if *W* has no inner join. Let *w*_*j*_ be its last inner split, or *w*_1_ if *W* has no inner split. The *core* of a walk is its subwalk from *w*_*j*_ to *w*_*i*_ if *j < i*, and walks with *i≥ j* have no core. A *simple omnitig* is a walk *W* that has a core^2^. The *univocal extension* of a walk *W* to be the maximal walk constructed by iteratively adding the unique out-neighbour of tail(*W*) to the end of *W* and iteratively adding the unique in-neighbour of head(*W*) to the beginning of *W*.

The *k-spectrum S*_*k*_ of a set of strings *S* is its set of substrings of length *k*. The (arc-centric) *de Bruijn graph* of a *k*-spectrum *S*_*k*_ is defined by vertex set *S*_*k−*1_ and for each *x ∈ S*_*k*_, an edge from the *k−* 1 prefix of *x* to the *k−* 1 suffix of *x*. In a de Bruijn graph, each walk *spells* a string by spelling out its first node, and then appending the last character of each subsequent node in order.

### 3.2 Safety of simple omnitigs

Informally, a walk in a genome graph is *safe* if it is guaranteed to be in any genome that could have generated the genome graph. Here we will focus on the arc centric de Bruijn graph of the reads as the genome graph. We cannot hope to generate only safe contigs if sequencing errors remain in the graph, so we assume, as in previous work, that all errors have been corrected [TM17]. A recent work also showed that if there are gaps in coverage, then even the unitig algorithm is not safe [RM22]. We therefore assume in our theoretical model that we have error-free reads with perfect coverage, and so our genome graph is the de Bruijn graph built on the *k*-spectrum of the genome.

In this setting, it is already known that simple omnitigs are guaranteed to be substrings of a singe-chromosome circular genome [Cai+20]. However, linear multi-chromosome genomes pose a challenge; for example, no safe and complete algorithm is known in this setting. However, as the following Theorem shows, the conditions for a simple omnitig to not be substring in this setting are very narrow.

#### Theorem 1.

*Let k* ∈ ℕ *and let S*_*k*_ *be the error-free k-spectrum of a linear genome with multiple chromosomes. Let G* = (*V, E*) *be the arc-centric de Bruijn graph of S*_*k*_. *Let L be the set of k−* 1*-mers that are the first or last k-mer of some chromosome. If none of the inner nodes of a simple omnitig are in L, then its spelled string is a substring of some chromosome of the genome*.

*Proof*. Let *W* ^*′*^ = (*w*_1_, …, *w*_*j*_, …, *w*_*i*_, … *w*_*ℓ*_) be a simple omnitig, where *W* = (*w*_*j*_, …, *w*_*i*_) is its core with *j < i* by definition. By definition, all nodes *w*_2_, …, *w*_*i−*1_ have a single incoming arc, and all nodes *w*_*j*+1_, …, *w*_*ℓ−*1_ have a single outgoing arc. Hence, for a walk that does not start or end with any inner node of *W* ^*′*^ to contain the arc (*w*_*j*_, *w*_*j*+1_), it needs to contain *W* ^*′*^ as subwalk.

Since each arc in *E* represents a *k*-mer of the genome, (*w*_*j*_, *w*_*j*+1_) is part of the genome, so it is contained in some chromosome. Each chromosome is an *s*-*t* walk *C* in *G* where *s, t* ∈ *L*, so there is some *C* that contains (*w*_*j*_, *w*_*j*+1_). By the argument above, this means that it contains *W* ^*′*^ as subwalk, so since *G* is error-free, *W* ^*′*^ is substring of the original genome.

Thus, simple omnitigs are safe in the case of multiple linear chromosomes, as long as they do not contain the start or end *k*-mer of a chromosome. Note that these conditions are not complete, since it is known [Cai+20] that there are also simple omnitigs containing ends of chromosomes that are safe, based on more complex conditions about the topology of the graph.

In practice, missing coverage and errors in the reads may still cause simple omnitigs to contain more misassemblies than unitigs, even though both are safe in theory. Missing coverage or errors may cause branching nodes to miss some branches, allowing simple omnitigs to extend over a branching node where a unitig would stop. For example, if a node has two incoming arcs and one outgoing arc, then a unitig would stop there, while a simple omnitig may extend over the node. However, if the node is actually missing a second outgoing arc due to missing coverage or errors, then the simple omnitig would not be safe in the error-free graph that includes the missing branch. Hence it possibly has a misassembly at the branching node.

### 3.3 Computing simple omnitigs

In this section, we give an algorithm to compute simple omnitigs in any graph in linear output-sensitive time. Note that we cannot simply output all univocal extensions of maximal unitigs, because that would generate simple omnitigs that are non-maximal (i.e. a simple omnitig may have more than one unitig as a subwalk). Instead, our algorithm iterates over all maximal unitigs and checks that 1) if the first node has exactly one outgoing arc then it has no incoming arcs, and that 2) if the last node has exactly one incoming arc, then it has no outgoing arc. As we prove below, these two conditions hold if and only if the unitig is a core of some maximal simple omnitig. For those unitigs where these conditions hold, the algorithm outputs their univocal extension as a simple omnitig. The correctness of the algorithm follows from the following theorem:

#### Theorem 2.

*The* core of a maximal simple omnitig *is a walk W* = (*w*_1_, …, *w*_*ℓ*_) *with ℓ*≥1 *such that*

*(a) the core of W is W, and*

*(b) if w*_1_ *has exactly one outgoing arc, then it has no incoming arcs, and*

*(c) if w*_*ℓ*_ *has exactly one incoming arc, then it has no outgoing arcs*.

*Proof:* Note that the cores of maximal simple omnitigs are unitigs by definition. (*⇒*) Let *W* ^*′*^ be the univocal extension of *W*, and a maximal simple omnitig. Then by definition, *W* is its core. Further, by definition, the core of a core *W* is *W* itself, so (a) holds. Next, if *w*_1_ has exactly one outgoing arc, then, since *W* ^*′*^ is maximal, *w*_1_ cannot have exactly one incoming arc. If it had more than one incoming arc, then *W* ^*′*^ would start at *w*_1_ and the univocal extension of any incoming arc would contain *W*, and hence the whole *W* ^*′*^. Since it is a univocal extension, it is a simple omnitig, so *W* ^*′*^ would not be maximal. Hence, if *w*_1_ has exactly one outgoing arc, then it has no incoming arcs, so (b) holds. Symmetrically, if *w*_*ℓ*_ has exactly one incoming arc, then it has no outgoing arcs, so (c) holds.

(*⇐*) Assume that *W* is not the core of a maximal simple omnitig. Then either (a) does not hold, or *W* is the core of a non-maximal simple omnitig, in which case its univocal extension *W* ^*′*^ is the subwalk of a maximal simple omnitig *X*^*′*^ with a core *X* ≠ *W*. Also, *W* is inside the univocal extension of *X*, and they share at most one node, which is the first of one and the last of the other. If *W* is right of *X* in *X*^*′*^, then *w*_1_ has exactly one outgoing arc, but at least one incoming arc, so (b) does not hold. Symmetrically, if *W* is left of *X* in *X*^*′*^, then *w*_*ℓ*_ has exactly one incoming arc, but at least one outgoing arc, so (c) does not hold.

Iterating all maximal unitigs takes *O*(*m*) time, and there are at most *O*(*m*) maximal unitigs (not necessarily a tight bound). The check if a maximal unitig is the core of a maximal simple omnitig takes constant time, and computing and outputting the univocal extension takes time linear in the length of the univocal extension. Hence, computing all maximal simple omnitigs takes *O*(*m* + *out*) time, where *out* is the total length of the maximal simple omnitigs in the graph.

The algorithm is implemented in Rust in [Sch22c]. The code is in the subfolder `implementation` and can be run with `cargo run --release --` ^3^`compute-trivial-omnitigs --non-scc` (plus arguments specifying input and output files and formats). Note that since the assemblers use bidirected graphs, we first convert the bidirected graphs to doubled graphs as in [RM22].

### 3.4 Modifying QUAST-LG

We evaluate the assembly using QUAST-LG 5.0.2 [Mik+18]. Since QUAST uses minimap2 [Li18] for alignment which was reported in [Ban+22] to work better on homopolymer-compressed data, we run QUAST in homopolymer-compressed space by passing it a homopolymer-compressed reference and homopolymer-compressed contigs. When using simple omnitigs, contigs may overlap even when reported from a perfectly correct genome graph. Hence, none of the metrics that QUAST uses by default is able to accurately capture the contiguity of the assembly. Even the most advanced metric built into QUAST, the NGAx group of metrics (e.g. NGA50 or NGA75), produce higher numbers than they should if an assembler e.g. outputs the same contigs twice or outputs overlapping contigs.

We instead implement the EA*x*max group of metrics within QUAST. It is inspired by the E-size [Sal+12] which gives the expected value of the average length of a contig aligned to a uniformly randomly chosen base in the reference. The E-size has some weaknesses though. First, it is not robust against misassemblies. This is easily fixed in the same way as it is done for NGAx, by aligning the contigs to the reference and using continuous (enough) alignments to compute the metric. Second, the E-size uses averaging for each base, resulting in unwanted effects such as a reduced E-size for assemblers that report many short contigs that potentially overlap with others. This might not actually have been intended by the designers of the E-size, which wanted it to answer the question: “How many genes will be completely contained within assembled contigs or scaffolds, rather than split into multiple pieces?” [Sal+12]. A gene that is completely contained inside a long contig is not affected by also being contained partially within a short contig. So we choose to compute the maximum alignment to each reference base, instead of the average. Lastly, most works in genome assembly [RL20; Kol+19; Ban+22; Che+21; Nur+20] use percentile-based metrics like N50, NG50 or NGA50 and do not compute the expected value over a uniform distribution of the reference bases. To stick with existing literature, we therefore compute the EA*x*max metrics percentile-based as well.

We define the EA*x*max metric on the alignments as follows:

a. For each reference base, store the length of the maximum alignment to the reference base.
b. Sort the reference bases by length of maximum alignment descending, and report the value at index *x/*100 `*` *G*, where *G* is the length of the reference.

We additionally modified QUAST to output misassemblies that occur at the same position on the reference just once. Otherwise, since a single unitig could be output multiple times as part of different simple omnitigs, we would count the same misassembly on the same unitig multiple times. But a misassembly within a unitig is not introduced by reporting simple omnitigs, but instead an error produced by the assembler itself, so we only want to count it once. To compute the unique misassemblies, we first sort all misassemblies by their position on the reference, and then merge those that are less than X bp apart, however keeping the maximum length of a unique misassembly below X (instead of transitively merging misassemblies into one big unique one much longer than X). X is given by the extensive misassembly threshold in QUAST, which is 3000 by default for large genomes.

The modified version of QUAST is available at [Sch22d]. We run it with the following parameters: `--large -t 28 --min-alignment 20000 --extensive-mis-size 500000 --min-identity 90` (plus arguments specifying input and output files and formats). Increasing the minimum size of extensive misassemblies to 500000 is inspired by [Nur+20], which employ the same parameters to evaluate HiCanu on the same D. melanogaster data to avoid falsely reported misassemblies due to heterozygosity. We use the same parameters for C. elegans, to avoid misassemblies for the same reason, and additionally because our C. elegans individuum does not exactly match the reference.

### 3.5 Data availability and reproducibility

Our experiments are implemented with snakemake [Möl+21] and available here [Sch22c]. The repository contains a conda `environment.yml` file at the root, and in this conda environment the experiments can be reproduced by running `snakemake --cores 28 report all`. We have run our experiments on a cluster running slurm [YJG03].

For D. melanogaster, we use the same data as in [Nur+20], using the reference with the accession `GCF 000001215.4` and HiFi reads from the sequence read archive with the accession `SRR10238607` with a median and mean read length of 24.4kbp. We filtered the reference to include only assigned sequences (keep only Chr 2L, Chr 2R, Chr 3L, Chr 3R, Chr 4, Chr M, Chr X and Chr Y).

For C. elegans, we use the official reference of the C. elegans Sequencing Consortium [Seq98], available with accession `GCA 000002985.3`. We use HiFi reads submitted by the Korea Research Institute of Bioscience and Biotechnology with accession `SRR22137522`. The reference includes all autosomes I to V and one sex chromosome X.

## 4 Conclusions

Despite much work on both the theoretical and practical aspects of genome assembly, it has remained challenging to apply novel theoretical ideas to the practical setting. Omnitigs are a powerful construct within the safe-and-complete theoretical framework, however, due to their complexity, they have not been applied in practice. Instead, we have taken a simpler construct, called simple omnitigs, and shown that it both has provable theoretical guarantees and is amenable to being plugged into existing assemblers. By combining co-located unitigs, simple omnitigs are able to provide correct flanking context around repeats and variations.

On the theoretical side, we have shown that given a multi-chromosomal linear genome, error-free reads, and perfect coverage, simple omnitigs are safe except for some corner cases; in other words, they are guaranteed to be substrings of the original genome. Note that both the perfect coverage and error-free requirements are necessary to prove correctness even in the simpler case of unitigs [RM22].

On the practical side, we have injected simple omnitigs into two popular assemblers (wtdbg2 and Flye) and tested them on two HiFi datasets. On D. melanogaster, this gave substantial improvements in alignment-based assembly contiguity. In particular, our modifications to wtdbg2 improved the EA*x*max metrics to the point where they were higher than those of the previously best assembler HiCanu, while being more than 20 times faster than HiCanu. On C. elegans, we saw similar contiguity improvements, with simple omnitigs improving the EA*x*max metrics of both wtdbg2 and Flye; compared with other tested assemblers, the highest EA50max was due to the modified Flye assembler.

The above improvements come at the cost of a small increase in the number of misassemblies. To completely prevent simple omnitigs from introducing misassemblies, the assemblers seem to require additional modifications before integrating simple omnitigs. Assembler developers usually test the accuracy of the genome graph relying on unitigs and QUAST-LG metrics. However, by outputting simple omnitigs, we are uncovering other errors in the graph, including in the topology that connects between unitigs. Fixing those requires deeper understanding the internals of the assemblers and, perhaps, introducing simple omnitigs at an earlier stage of the development process. Another cause of misassemblies could be that gaps in coverage (which are not covered by the theory) destroy the correctness of some simple omnitigs (as they do for unitigs [RM22]). It would be interesting to understand the extent of this possible effect.

Though more work is needed to incorporate simple omnitigs into “production-ready” assemblers, our work overcomes many of the barriers that have held back omnitigs from being used in practice. First, omnitigs require complicated data structures and algorithms, while our algorithm to compute simple omnitigs is simple to implement and understand. Second, the theory of omnitig safety does not extend to the linear multichromosome setting, while we showed that, except for corner cases, simple omnitigs remain safe. Third, the omnitig algorithm is too slow to complete in a multi-chromosome setting, while the simple omnitig algorithm is fast. In conclusion, we hope this work helps to bridge the gap between theory and practice of genome assembly by adapting a complicated theoretical construct (i.e. omnitigs) to work in a practical setting.

## Acknowledgements

PM would like to thank John Hutton for early attempts to extend omnitigs to work in practice [Hut18]. The authors wish to thank the Finnish Computing Competence Infrastructure (FCCI) for supporting this project with computational and data storage resources. This work was partially funded by the European Research Council (ERC) under the European Union’s Horizon 2020 research and innovation programme (grant agreement No. 851093, SAFEBIO), and partially by the Academy of Finland (grants No. 322595, 328877). This material is based upon work supported by the National Science Foundation under Grants No. DBI-2138585, IIS 1453527 and OAC-1931531. Research reported in this publication was supported by the National Institute Of General Medical Sciences of the National Institutes of Health under Award Number R01GM146462. The content is solely the responsibility of the authors and does not necessarily represent the official views of the National Institutes of Health.

## A Omitted implementation details

### A.1 Modifying wtdbg2

The assembler wtdbg2 [RL20] comes with two binaries, the first `wtdbg2` which computes contigs as an overlapping sequence of reads, and the second `wtpoa-cns` which computes a consensus sequence for each of these contigs. All our modification happen in and around the first binary. The `wtdbg2` binary first builds a fuzzy de Bruijn graph and from which it computes unitigs after some error corrections, and then builds a fragment graph from these unitigs. In the fragment graph the assembler does another round of error corrections before it reports unitigs from this fragment graph and outputs their overlapping sequences of reads. We modified `wtdbg2` to output simple omnitigs instead of unitigs from the error corrected fragment graph. For this we run `wtdbg2` once and output the fragment graph using its builtin output functions. Then we compute simple omnitigs on this graph and run `wtdbg2` again, this time loading the simple omnitigs from disk and outputting their sequences of overlapping reads instead of the unitigs in the end. We can run `wtdbg2` two times this way and get consistent node/edge identifiers since these are deterministic, even when running `wtdbg2` in parallel.

In our experiments, we also realised that since HiFi reads have mostly base duplications and deletions as errors, we can homopolymer-compress the reads to get a more accurate assembly. It turned out that when reporting simple omnitigs, we get a lot more misassemblies, but when using homopolymer compression as well, the misassemblies are not a problem, at least for D. melanogaster. To still get an assembly in uncompressed space, we decompress the contigs before calling `wtpoa-cns`. This is simple since the overlapping sequences of reads that form the contigs of `wtdbg2` carry the ids of the original reads, so we can directly replace the subsequence of characters (adjusting the start and end index according to the compression). Note that, even though we evaluate in homopolymer compressed space in the end, this decompression is still important as it makes the consensus stage run in homopolymer decompressed space, thus producing an assembly in decompressed space. This allows for a fairer comparison of the modified wtdbg2 to the other assemblers, which all output contigs in uncompressed space. We call the variant of wtdbg2 that just uses homopolymer compression “W-int” and the variant that uses both homopolymer compression and simple omnitigs “W-so”. The modified wtdbg2 is available at [Sch22e]. Homopolymer compression is done with [Sch22b] and decompression is done with [Sch22f]. Hompolymer compression and decompression is implemented in Rust and run with `cargo run --release -- --compute-threads 28` (plus arguments specifying input and output files and formats). The modified wtdbg2 is run with the parameters proposed for HiFi reads: `-x ccs –g <REFERENCE LENGTH> -t 28` (plus arguments specifying input and output files and formats).

### A.2 Modifying Flye

The assembler Flye [Kol+19] runs in multiple stages. We have disabled resolving repeats and polishing, which does not seem to make any difference in our experiments. Instead, we directly report the unitigs of the repeat graph as computed by Flye. This variant is called “F-int”. When using simple omnitigs, we report simple omnitigs from the repeat graph instead of unitigs by calling our tool from within Flye (and we keep resolving repeats and polishing disabled). This variant is called “F-so”. The modified version of Flye is available at [Sch22a]. We run it with parameters `-g <REFERENCE LENGTH> -t 28 --pacbio-hifi` (plus arguments specifying input and output files and formats). To disable resolving of repeats and polishing, we add `--stop-after contigger`.

## Footnotes

1 This is the definition that is consistently used throughout the safe-and-complete literature (e.g. [Cai+20]), but we note that it is slightly different from one used in graph compaction literature (e.g. [CLM16]).

2 Note that this definition of simple omnitigs is equivalent to that in Section 2.1, except that it also allows subwalks of simple omnitigs to be simple omnitigs. This is to make our theoretical results more general, but in practice we only use maximal simple omnitigs.

3 This space is on purpose.

